# A universal chromosome identification system for maize and wild *Zea* species

**DOI:** 10.1101/2020.01.22.915942

**Authors:** Guilherme T. Braz, Lívia do Vale Martins, Tao Zhang, Patrice S. Albert, James A. Birchler, Jiming Jiang

**Affiliations:** Department of Plant Biology, Michigan State University, East Lansing, MI 48824, USA; Jiangsu Key Laboratory of Crop Genomics and Molecular Breeding/Key Laboratory of Plant Functional Genomics of Ministry of Education, Yangzhou University, 225009, China; Division of Biological Sciences, University of Missouri, Columbia, MO 65211, USA; Department of Horticulture, Michigan State University, East Lansing, MI 48824, USA; Michigan State University AgBioResearch, East Lansing, MI 48824, USA

**Keywords:** FISH, oligo-FISH, chromosome identification, karyotype, maize

## Abstract

Maize was one of the first eukaryotic species in which individual chromosomes can be identified cytologically, which made maize one of the oldest models for genetics and cytogenetics research. Nevertheless, consistent identification of all 10 chromosomes from different maize lines as well as from wild *Zea* species remains a challenge. We developed a new technique for maize chromosome identification based on fluorescence in situ hybridization (FISH). We developed two oligonucleotide-based probes that hybridize to 24 chromosomal regions. Individual maize chromosomes show distinct FISH signal patterns, which allow universal identification of all chromosomes from different *Zea* species. We developed karyotypes from three *Zea mays* subspecies and two additional wild *Zea* species based on individually identified chromosomes. A paracentric inversion was discovered on the long arm of chromosome 4 in *Z. nicaraguensis* and *Z. luxurians* based on modifications of the FISH signal patterns. Chromosomes from these two species also showed distinct distribution patterns of terminal knobs compared to other *Zea* species. These results support that *Z. nicaraguensis* and *Z. luxurians* are closely related species.

## Introduction

A reliable and efficient system for chromosome identification is the foundation for cytogenetic research. Maize (*Zea mays*) and common fruit fly (*Drosophila melanogaster*) became the most important model species for genetic and cytogenetic research in early 20^th^ century because individual chromosomes could be readily identified in both species (Bridges 1935; McClintock 1929). However, the identification of individual chromosomes relied on meiotic pachytene chromosomes in maize and salivary polytene chromosomes in *Drosophila*. Thus, the chromosome identification methodologies developed in these two model species were not applicable in most of the other eukaryotes. Preparation and identification of pachytene chromosomes is technically demanding, thus, cytogenetic research in most plant species relies on mitotic metaphase chromosomes. Most of the chromosome identification systems based on mitotic metaphase chromosomes were developed after 1970s using either chromosome banding or DNA in situ hybridization techniques (Jiang and Gill 1994).

Although maize chromosomes can be individually identified based on their morphology (McClintock 1929), there are ambiguities. Chromosome banding techniques were developed in 1980s and 1990s for maize chromosome identification (de Carvalho and Saraiva 1993, 1997; Deaguiarperecin and Vosa 1985; Jewell and Islam-Faridi 1994; Kakeda et al. 1990; Rayburn et al. 1985; Ward 1980). C-banding, the most popular banding technique in plants, revealed mostly the knob-related regions on maize chromosomes (Deaguiarperecin and Vosa 1985; Jewell and Islam-Faridi 1994; Rayburn et al. 1985). Maize chromosomes showed a limited number of C-bands that are also highly polymorphic among different maize varieties. Therefore, a standard banding pattern cannot be established for routine maize chromosome identification.

Fluorescence in situ hybridization (FISH) gradually replaced chromosome banding to become the most popular technique for chromosome identification in plants (Jiang and Gill 2006). Repetitive DNA sequences and large-insert DNA clones, such as bacterial artificial chromosome (BAC) clones, have been most commonly used as FISH probes in maize (Amarillo and Bass 2007; Chen et al. 2000; Figueroa and Bass 2012; Koumbaris and Bass 2003; Lamb et al. 2007b; Sadder and Weber 2001; Wang et al. 2006). Kato et al. (2004) used a combination of several repetitive DNA sequences as FISH probes to identify all 10 maize chromosomes in the same metaphase cells. This repeat-based FISH method was successfully used to identify chromosomes in various maize lines as well as wild *Zea* species and subspecies (Albert et al. 2010; Kato et al. 2004). The FISH signal patterns of some repeats, however, are highly polymorphic among different maize lines, which may cause difficulties or errors in chromosome identification.

Recently, a new class of FISH probes was developed based on pooled single-copy oligonucleotides (oligo) (Jiang 2019). If the genomic sequence is available from a target species, oligos, typically 40-50 bp long, specific to a chromosomal region or to an entire chromosome can be synthesized in parallel and labeled as a FISH probe (Han et al. 2015). Oligo-FISH probes have been successfully developed in several plant species for various types of cytogenetic studies (Albert et al. 2019; Bi et al. 2020; He et al. 2018; Hou et al. 2018; Martins et al. 2019; Meng et al. 2018; Qu et al. 2017; Simonikova et al. 2019; Song et al. 2020; Xin et al. 2018; Xin et al. 2020). A FISH probe can include oligos derived from multiple regions from multiple chromosomes (Braz et al. 2018). Such oligo-FISH probes generate specific hybridization patterns on individual chromosomes, which resemble distinct barcodes on different chromosomes. This chromosome identification strategy has been successfully demonstrated in potato (Braz et al. 2018), rice (Liu et al. 2020) and sugarcane (Meng et al. 2020). Here, we report the development of similar barcode FISH probe in maize. We demonstrate that barcode FISH can be used as an universal chromosome identification system for maize and all wild *Zea* species. The FISH signal pattern is not affected by the DNA polymorphisms existing among different maize varieties or even among different species. Using this system we were able to establish karyotypes of several different *Zea* species and subspecies and to discover a paracentric inversion specific to two wild *Zea* species.

## Materials and Methods

### Plant Materials

We included the following *Zea* species and subspecies in chromosome identification and karyotyping studies: *Z. mays* (inbred B73), *Z. mays* ssp. *parviglumis* (Ames 21826), *Z. mays* ssp. *mexicana* (Ames 21851), *Z. mays* ssp. *huehuetenangensis* (PI 441934), *Z. diploperennis* (PI 462368), *Z. nicaraguensis* (PI 615697), and *Z. luxurians* (PI 422162). Additional species included in the study were the tetraploid species *Z. perennis* (Ames 21874), *Tripsacum dactyloides* (PI 421612), and *Sorghum bicolor* (PI 564163). Seeds of all wild species were originally from the Germplasm Resources Information Network (GRIN) and the National Genetics Resource Program (Ames, Iowa).

### Oligo-FISH probe design

The two oligo-based barcode probes were designed following our published procedure (Albert et al. 2019) using Chorus software (https://github.com/forrestzhang/Chorus). Briefly, the maize genome sequence was divided into 45-nt long oligos in step size of 3 nt. The short sequence reads were mapped back to the genome using BWA (Burrows-Wheeler Alignment tool) (Li and Durbin 2009). Oligos mapped to two or more locations (with 70% homology) were eliminated. All repetitive sequences related oligos were filtered out from the oligo set using *k*-mer method (Albert et al. 2019). We selected oligos from 24 chromosomal regions to create a barcode for all 10 maize chromosomes (**Figure 1a**). Each selected region contains approximately 2000 oligos. We selected the chromosomal regions that are relatively enriched with single copy sequences. Each of the 24 selected regions spans 1.3-3.3 Mb (**Table S1**). If two regions are designed from the same chromosomal arm, we then intended to separate the two regions as far as possible. The two regions on eight different chromosomal arms (**Figure 1a**) are separated 28.2-66.4 Mb (**Table S1**).

**Figure 1.**
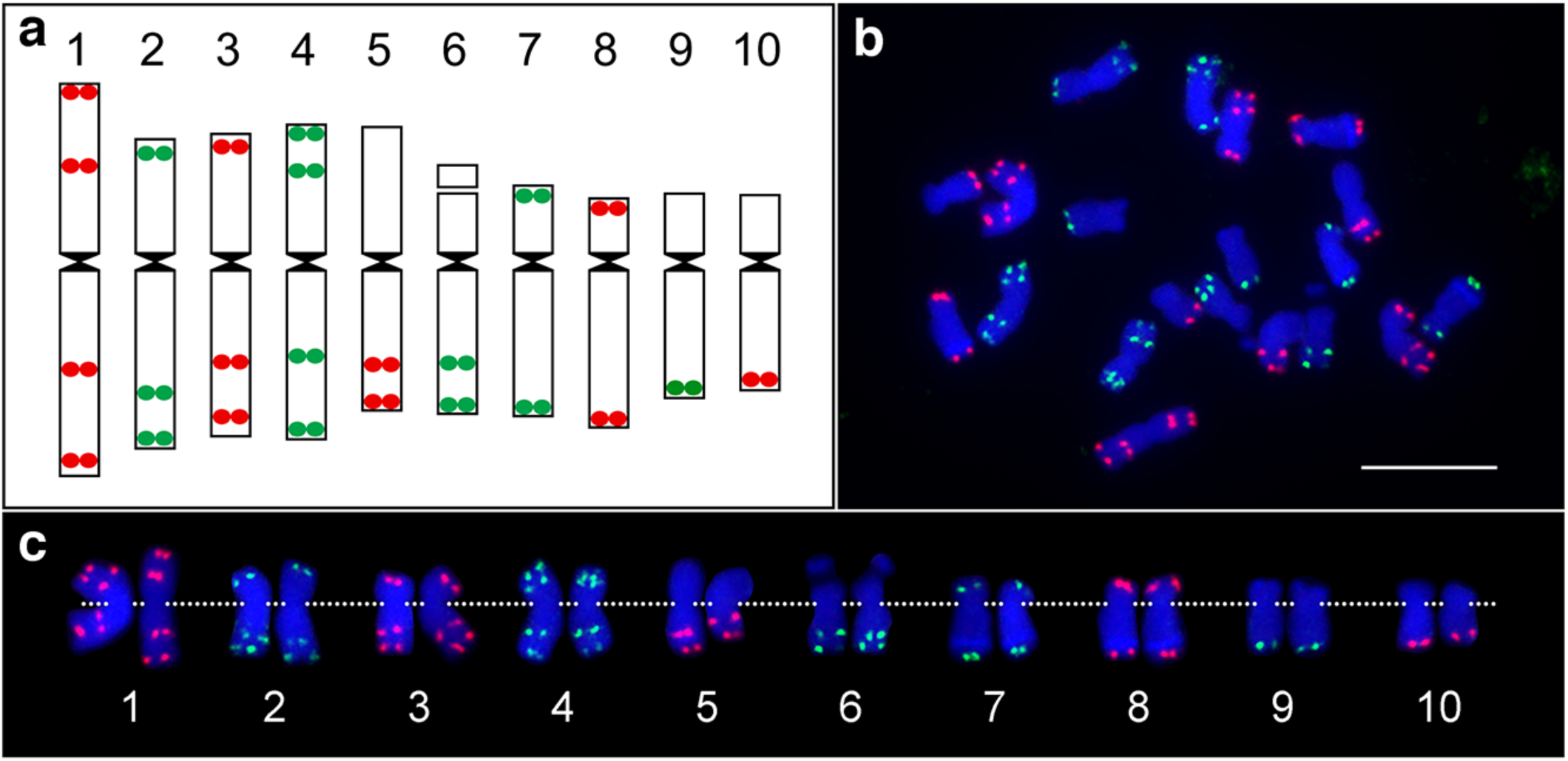
Development of barcode oligo-FISH probes for maize chromosome identification. **(a)** Predicted locations of the oligo-FISH signals on 10 maize chromosomes. Oligos were selected from a total of 24 chromosomal regions (12 red regions and 12 green regions). The 10 chromosomes can be distinguished from each other based on number and location of the red/green signals. The centromere positions on the 10 chromosomes in the maize reference genome were based on the locations of sequences associated with CENH3 nucleosomes (Zhao et al., 2016). **(b)** FISH mapping of the two oligo-FISH probes on metaphase chromosomes prepared from maize inbred B73. Bar = 10 µm. **(c)** Homologous chromosomes were digitally excised from (b) and paired. The centromeres of the chromosomes are aligned by a dotted line.

### Oligo-FISH

Root tips harvested from plants grown in greenhouse were treated with nitrous oxide at a pressure of 160 psi (∼10.9 atm) for 2 hrs. and 20 min (Kato 1999), fixed in fixative solution (3 ethanol : 1 acetic acid) and kept at −20°C. Root tips were digested using an enzymatic solution composed by 4% cellulase (Yakult Pharmaceutical, Japan), 2% pectinase (Sigma-Aldrich Co., USA) and 2% pectolyase (Plant Media, USA) for two hours at 37°C and slides were prepared using the stirring method (Ross et al. 1996).

Two oligo-based FISH probes were labeled according to published protocols (Han et al. 2015) and hybridized to metaphase chromosomes (Cheng et al. 2002). Biotin- and digoxygenin-labeled probes were detected by anti-biotin fluorescein (Vector Laboratories, Burlingame, California) and anti-digoxygenin rhodamine (Roche Diagnostics, Indianapolis, Indiana), respectively. Chromosomes were counterstained with 4,6-Diamidino-2-phenylindole (DAPI) in VectaShield antifade solution (Vector Laboratories). FISH images were captured using a QImaging Retiga EXi Fast 1394 CCD camera attached to an Olympus BX51 epifluorescence microscope. Images were processed with Meta Imaging Series 7.5 software. The final contrast of the images was processed using Adobe Photoshop software.

### Karyotyping

We used 7-10 complete metaphase cells of *Z. mays* and its relatives (*Z. mays* ssp. *parviglumis, Z. mays* ssp. *mexicana, Z. mays* ssp. *huehuetenangensis, Z. nicaraguensis, Z. luxurians*) to measure the short (S) and long (L) arms of individual chromosomes using DRAWID software version 0.26 (Kirov et al. 2017). The measurements were used to determine the total length of each chromosome (*tl* = *S* + *L*), total length of entire set of chromosomes (*TL* = Σ*tl*), arm ratio (*AR* = *L*/*S*) of each chromosome, and relative length of each chromosome (*RL* = *tl*/*TL* × 100). Chromosomal knobs were identified as DAPI-positive bands. Chromosomes were classified based on arm ratio following Levan et al. (1964).

## Results

### Development of oligo-FISH probes for maize chromosome identification

We developed two oligo-based FISH probes to facilitate simultaneous identification of all 10 maize chromosomes in the same metaphase cells. These two probes contained a total of 50,082 oligos (45 nt) designed from single copy sequences in the maize genome (Schnable et al. 2009). The oligos were selected from 24 chromosomal regions (**Figure 1a**), each containing 1978-2282 oligos and spanning 1.3-3.3 megabase (Mb) of DNA sequences (**Table S1**). The oligos were synthesized as two separate pools, containing 25,059 (red probe) and 25,023 (green probe) oligos, respectively. FISH using these two probes generated 24 distinct signals on metaphase chromosomes prepared from maize inbred B73 (**Figure 1b**). The chromosomal positions of the FISH signals matched with predicted positions on the 10 pseudomolecules, which allowed us to readily distinguish 10 individual chromosomes in the same metaphase cell (**Figure 1c**). Several chromosomal arms contained two signals, which were well separated on metaphase chromosomes. The two signals designed on the short arm of chromosome 4 were separated by 28.2 Mb, representing the shortest distance among the paired signals (**Table S1**). These two signals appeared to be fused only on some highly condensed metaphase chromosomes.

### Chromosome identification in different *Zea* species

The two oligo-FISH probes were used to identify chromosomes from several *Z. mays* subspecies and wild *Zea* species known as “teosintes”, including *Z. mays* ssp. *parviglumis* (**Figure 2a**), *Z. mays* ssp. *mexicana* (**Figure 2b**), *Z. mays* ssp. *huehuetenangensis* (**Figure 2c**), *Z. diploperennis* (**Figure 2d**), *Z. nicaraguensis* (**Figure 2e**), and *Z. luxurians* (**Figure 2f**). We observed a nearly identical FISH signal pattern on chromosomes from all *Zea* species and subspecies (**Figure 3**). We also performed oligo-FISH analysis in the tetraploid species *Z. perennis* (2n = 4x = 40). Similarly, the conserved FISH signal pattern was observed on all 10 sets of homologous chromosomes (**Figure 4**). Thus, these two probes allowed us to identify all chromosomes from all *Zea* taxa.

**Figure 2.**
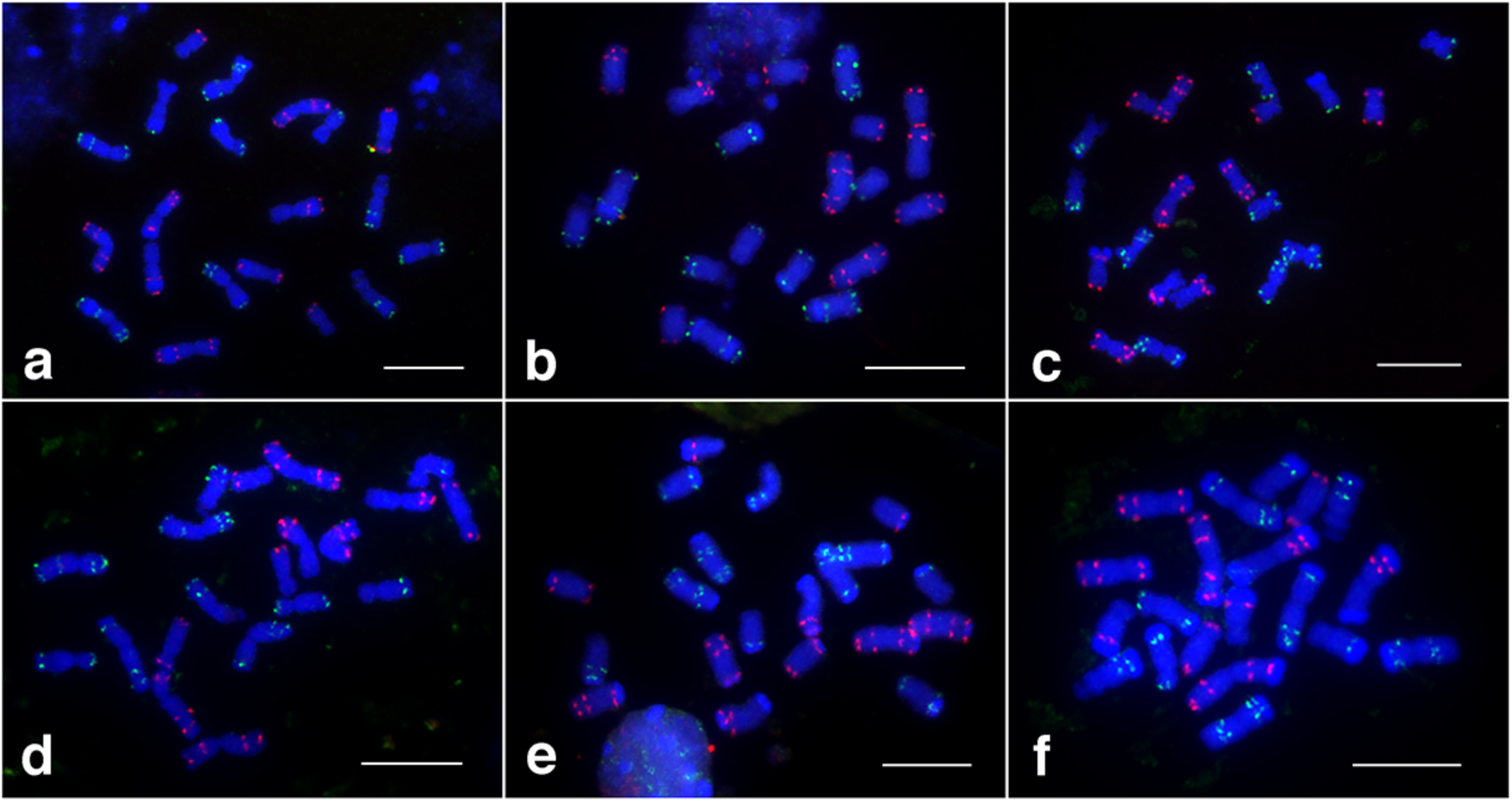
FISH mapping of the two oligo-FISH probes on metaphase chromosomes prepared from **(a)** *Z. mays* ssp. *parviglumis*; **(b)** *Z. mays* ssp. *mexicana*; **(c)** *Z. mays* ssp. *huehuetenangensis*; **(d)** *Z. diploperennis*; **(e)** *Z. nicaraguensis*; and **(f)** *Z. luxurians*. Bars = 10 µm.

**Figure 3.**
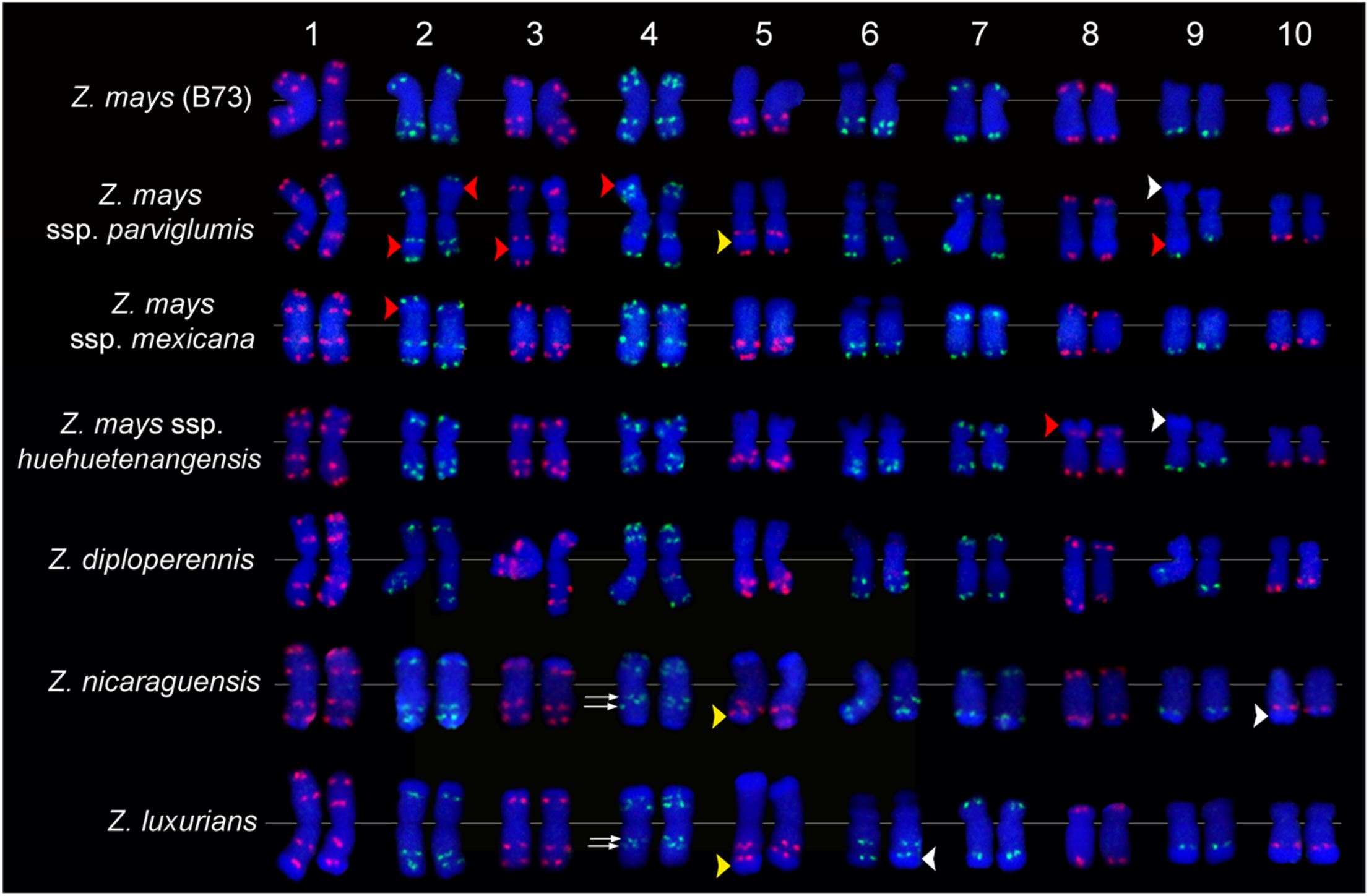
Comparative karyotyping of seven *Zea* species and subspecies. Chromosomes 1-10 from each species/subspecies are arranged from left to right. The karyotypes were developed from the same metaphase cells in Figure 1b for maize and Figure 2 for other species/subspecies. Double arrows point to the two green signals on the long arm of chromosome 4 of *Z. nicaraguensis* and *Z. luxurians*. Both signals are located in the interstitial regions. Yellow arrowheads point to a knob on chromosome 5, which is located in the subterminal region in *Z. mays* ssp. *parviglumis*, but at the distal end in *Z. nicaraguensis* and *Z. luxurians*. Red arrowheads point to knobs observed only on one of the two homologous chromosomes. White arrowheads point to knobs that show visibly different sizes on homologous chromosomes.

**Figure 4.**
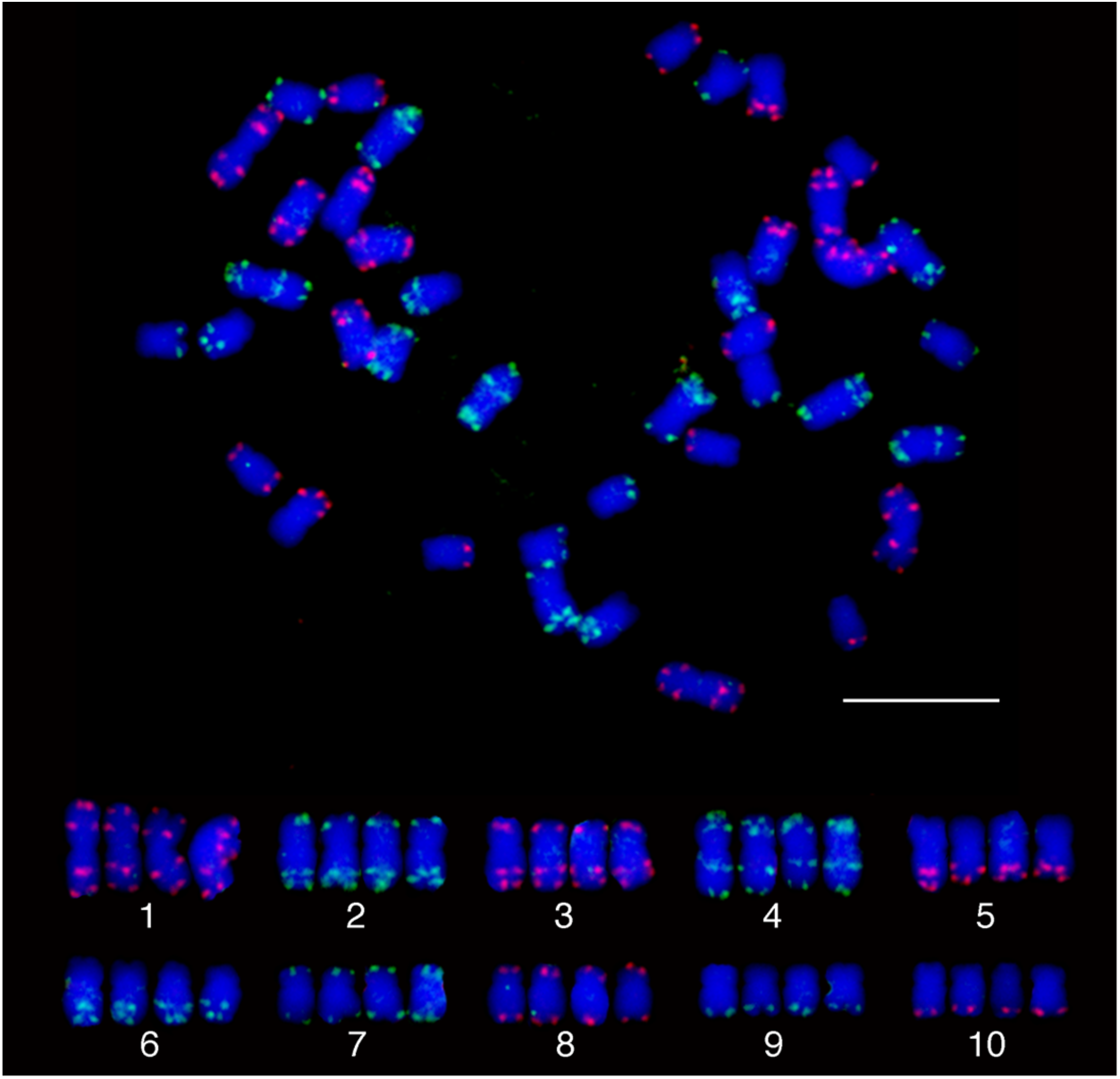
Chromosome identification in a tetraploid species *Z. perennis*. The top panel shows a complete metaphase cell hybridized with the two oligo-FISH probes. The bottom panel shows the 4 homologous chromosomes of each of the 10 *Z. perennis* chromosomes digitally excised from the same cell. Bar =10 µm.

### A putative inversion detected on long arm of chromosome 4

We detected some noticeable chromosome level differences based on the signal patterns of oligo-FISH. One distinct FISH signal pattern change was observed on the long arm of chromosome 4 from *Z. nicaraguensis* and *Z. luxurians* (**Figure 3**). One green signal was designed from the distal end on the long arm of chromosome 4 (at 234-236 Mb of chromosome 4, which is 242 Mb long). This signal was detected at the end of the long arm of maize chromosome 4 (**Figure 1c**). However, chromosome 4 of *Z. nicaraguensis* and *Z. luxurians* did not contain this terminal signal. It instead contains two interstitial green signals on the long arm (**Figure 3**). Thus, a paracentric inversion may have occurred in the long arm, which would relocate the terminal signal into an interstitial position.

### Chromosomal difference revealed by oligo-FISH

The number, size, and distribution of metaphase-visible “knobs” were the most prominent factor resulting in the difference observed on the same chromosome from different species (**Figure 3**). Although the polymorphism of knob distribution among different maize lines is well known (Adawy et al. 2004; Albert et al. 2010), distinct knob distribution patterns were observed in some species. For example, a knob was detected between the two red signals on the long arm of chromosome 5 in *Z. mays* ssp. *parviglumis*. By contrast, a knob was observed at the distal end of the long arm of chromosome 5 in *Z. nicaraguensis* and *Z. luxurians* (**Figure 3**, yellow arrowheads). A similar difference of FISH signal position among species, caused by the presence of a terminal vs. subterminal knob, was observed on the distal regions of several other chromosomal arms, including the short arm of chromosome 2, and long arm of chromosomes 1, 2, 3, 6, 7, 8, and 9 (**Figure 3**). Interestingly, most of the terminal knobs in *Z. nicaraguensis* and *Z. luxurians* were located distal to the FISH signals, suggesting a close structural similarity of these two species (**Figure 3**).

In several cases, a major knob was observed only on one copy of a pair of homologous chromosomes. For example, the long arm of chromosome 3 contained two red signals (**Figure 1a**). The two red signals were separated by a major knob in one copy of chromosome 3 in *Z. mays* ssp. *parviglumis*. This knob was not visible in the second copy of chromosome 3 (**Figure 3**). The distance between the two red signals appeared to be significantly different on the two chromosomes due to the presence of this heterozygous knob. A similar heterozygous knob was observed on several other chromosomes among different species (**Figure 3**, red arrowheads).

The size of a knob can also be visibly different on two homologous chromosomes. For example, a terminal knob was observed on the long arm of chromosome 10 in several species. The size of this knob appeared to be distinctly different on the two copies of chromosome 10 in *Z. nicaraguensis* (**Figure 3**, white arrowheads). Both presence/absence and size polymorphism of major knobs can result in a significant difference of the size of the homologous chromosomes.

### The karyotypes of different *Zea* species and subspecies

Unambiguous identification of all chromosomes in the same metaphase cells allowed us to develop karyotypes based on individually identified chromosomes from all *Zea* species and subspecies (**Tables 1, 2**). Each karyotype was developed based on measurements of all chromosomes in 7-10 complete metaphase cells. Not surprisingly, all *Zea* species and subspecies shared a highly similar karyotype with similar relative length and arm ratio of all 10 chromosomes (**Tables 1, 2**). Chromosomes 1, 4, 5, and 10 are morphologicaly conserved, and are metacentric (Chr. 1, 4, and 5) and submetacentric (Chr. 10) in all species (**Table 2**). The other chromosomes are also similar but with minor variations among different species. Chromosomes 2 and 3 are metacentric in all species, except in *Zea mays* ssp. *mexicana* (submetacentric). Chromosome 7 is submetacentric in all species, except *Z. mays* ssp. *mexicana* (metacentric). Chromosome 8 is submetacentric in all the species, except in *Z. mays* ssp. *huehuetenangensis* (metacentric). Chromosome 9 is submetacentric in all the species, except *Z. mays* ssp. *parviglumis* and *Z. mays* ssp. *huehuetenangensis* (metacentric). Chromosome 6 is the only satellite chromosomes in all species.

**Table 1.**
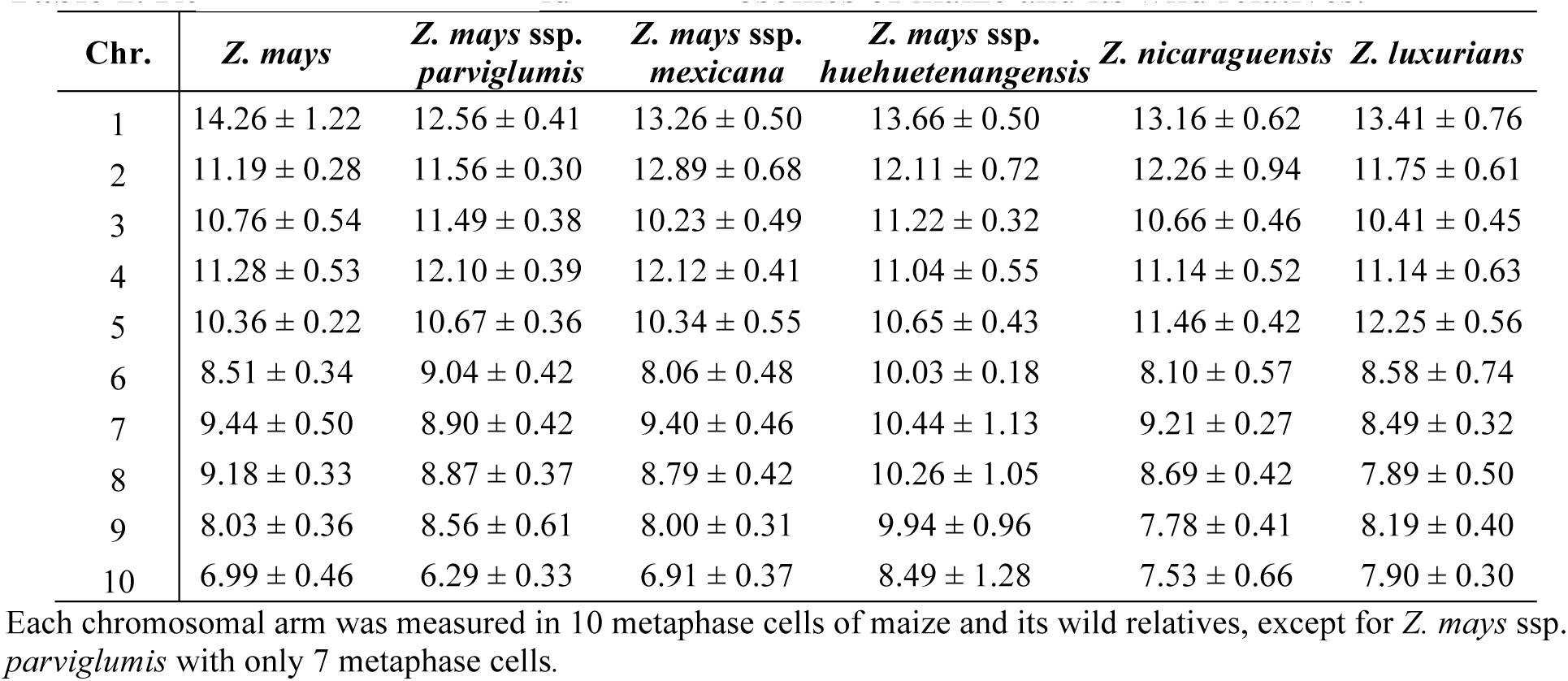
Relative length of individual chromosomes of maize and its wild relatives.

| Chr. | *Z. mays* | *Z. mays* ssp.<br>*parviglumis* | *Z. mays* ssp.<br>*mexicana* | *Z. mays* ssp.<br>*huehuetenangensis* | *Z. nicaraguensis* | *Z. luxurians* |
| --- | --- | --- | --- | --- | --- | --- |
| 1 | 14.26 ± 1.22 | 12.56 ± 0.41 | 13.26 ± 0.50 | 13.66 ± 0.50 | 13.16 ± 0.62 | 13.41 ± 0.76 |
| 2 | 11.19 ± 0.28 | 11.56 ± 0.30 | 12.89 ± 0.68 | 12.11 ± 0.72 | 12.26 ± 0.94 | 11.75 ± 0.61 |
| 3 | 10.76 ± 0.54 | 11.49 ± 0.38 | 10.23 ± 0.49 | 11.22 ± 0.32 | 10.66 ± 0.46 | 10.41 ± 0.45 |
| 4 | 11.28 ± 0.53 | 12.10 ± 0.39 | 12.12 ± 0.41 | 11.04 ± 0.55 | 11.14 ± 0.52 | 11.14 ± 0.63 |
| 5 | 10.36 ± 0.22 | 10.67 ± 0.36 | 10.34 ± 0.55 | 10.65 ± 0.43 | 11.46 ± 0.42 | 12.25 ± 0.56 |
| 6 | 8.51 ± 0.34 | 9.04 ± 0.42 | 8.06 ± 0.48 | 10.03 ± 0.18 | 8.10 ± 0.57 | 8.58 ± 0.74 |
| 7 | 9.44 ± 0.50 | 8.90 ± 0.42 | 9.40 ± 0.46 | 10.44 ± 1.13 | 9.21 ± 0.27 | 8.49 ± 0.32 |
| 8 | 9.18 ± 0.33 | 8.87 ± 0.37 | 8.79 ± 0.42 | 10.26 ± 1.05 | 8.69 ± 0.42 | 7.89 ± 0.50 |
| 9 | 8.03 ± 0.36 | 8.56 ± 0.61 | 8.00 ± 0.31 | 9.94 ± 0.96 | 7.78 ± 0.41 | 8.19 ± 0.40 |
| 10 | 6.99 ± 0.46 | 6.29 ± 0.33 | 6.91 ± 0.37 | 8.49 ± 1.28 | 7.53 ± 0.66 | 7.90 ± 0.30 |

Each chromosomal arm was measured in 10 metaphase cells of maize and its wild relatives, except for *Z. mays* ssp. *parviglumis* with only 7 metaphase cells.

**Table 2.** Arm ratio of individual chromosomes of maize and its wild relatives.

| Chr. | *Z. mays* | *Z. mays* ssp.<br>*parviglumis* | *Z. mays* ssp.<br>*mexicana* | *Z. mays* ssp.<br>*huehuetenangensis* | *Z. nicaraguensis* | *Z. luxurians* |
| --- | --- | --- | --- | --- | --- | --- |
| 1 | 1.26 ± 0.14 m | 1.22 ± 0.16 m | 1.25 ± 0.11 m | 1.26 ± 0.27 m | 1.11 ± 0.12 m | 1.16 ± 0.12 m |
| 2 | 1.52 ± 0.18 m | 1.49 ± 0.36 m | 1.83 ± 0.32 sm | 1.43 ± 0.21 m | 1.38 ± 0.23 m | 1.27 ± 0.17 m |
| 3 | 1.65 ± 0.28 m | 1.61 ± 0.27 m | 1.89 ± 0.34 sm | 1.53 ± 0.19 m | 1.44 ± 0.18 m | 1.59 ± 0.19 m |
| 4 | 1.34 ± 0.17 m | 1.51 ± 0.32 m | 1.60 ± 0.30 m | 1.26 ± 0.17 m | 1.55 ± 0.21 m | 1.23 ± 0.17 m |
| 5 | 1.54 ± 0.45 m | 1.59 ± 0.25 m | 1.68 ± 0.32 m | 1.31 ± 0.35 m | 1.34 ± 0.27 m | 1.16 ± 0.16 m |
| 6 | 2.40 ± 0.46 sm | 3.31 ± 0.90 a | 2.74 ± 0.63 sm | 2.39 ± 0.39 sm | 3.32 ± 1.06 a | 3.56 ± 0.60 a |
| 7 | 2.11 ± 0.46 sm | 2.53 ± 0.58 sm | 1.51 ± 0.37 m | 1.86 ± 0.22 sm | 2.58 ± 0.65 sm | 2.39 ± 0.35 sm |
| 8 | 2.13 ± 0.48 sm | 2.78 ± 0.75 sm | 1.97 ± 0.47 sm | 1.69 ± 0.33 m | 1.99 ± 0.48 sm | 2.29 ± 0.32 sm |
| 9 | 1.80 ± 0.34 sm | 1.58 ± 0.29 m | 1.88 ± 0.35 sm | 1.41 ± 0.34 m | 2.77 ± 0.58 sm | 2.89 ± 0.63 sm |
| 10 | 1.78 ± 0.40 sm | 1.99 ± 0.20 sm | 1.83 ± 0.37 sm | 1.99 ± 0.36 sm | 2.53 ± 0.38 sm | 2.81 ± 0.39 sm |

Each chromosomal arm was measured in 10 metaphase cells of maize and its wild relatives, except for *Z. mays* ssp. *parviglumis* with only 7 metaphase cells. m = metacentric, sm = submetacentric, a = acrocentric.

### Application of the oligo probes in distantly related species

We explored the potential of the barcode probes for chromosome identification in species that are more distantly related to maize. We first tested the two maize probes in hybridization to metaphase chromosomes prepared from *Tripsacum dactyloides* (2n = 2x = 36), a species from a sister genus related to *Zea* and diverged from maize about 4.5 million years (MYs) (Hilton and Gaut 1998). The two oligo-FISH probes produced punctuated signals on most of chromosomes, which allowed to identify few putative homoeologous chromosomes (**Figure 5a**). However, most of the signals were not as strong as those on maize chromosomes. In addition, strong backgorund signals were observed on most chromosomes. Several *T. dactyloides* chromosomes lacked unambiguous signals (**Figure 5a**). Thus, the maize oligo probes were not useful to identify individual *T. dactyloides* chromosomes.

**Figure 5.**
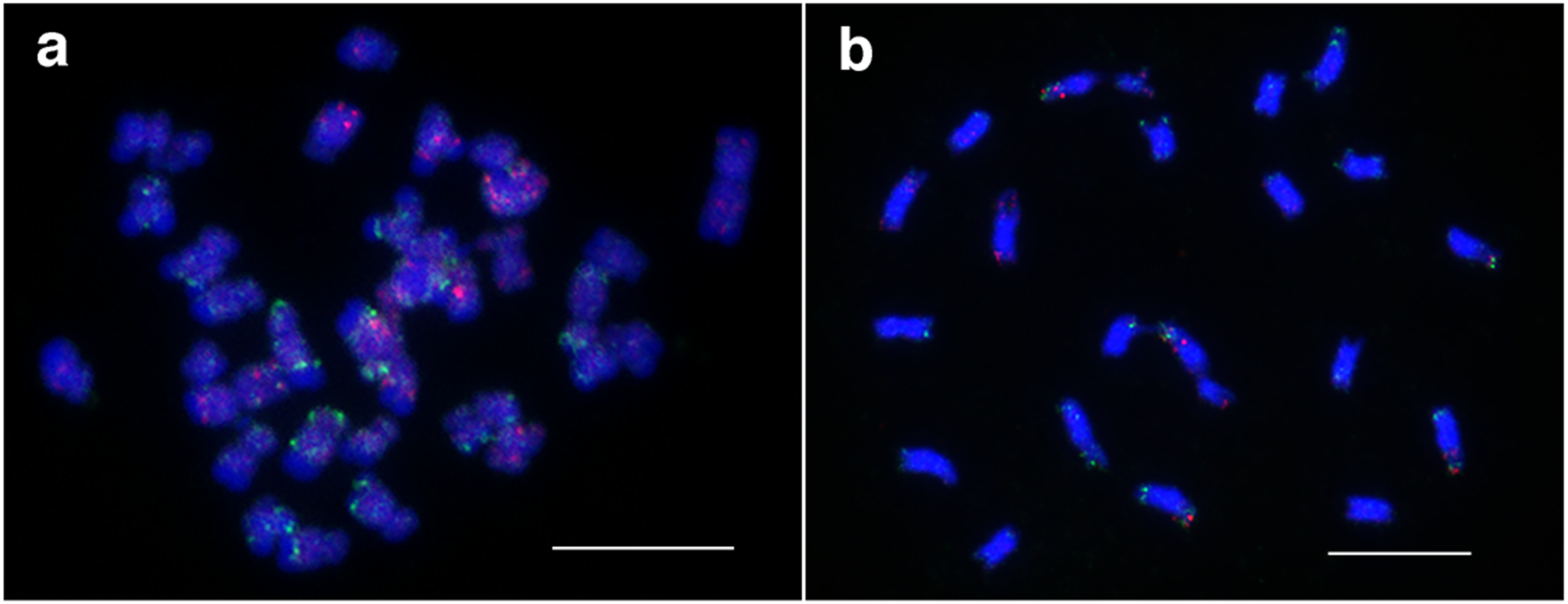
FISH mapping of the two oligo-FISH probes on metaphase chromosomes prepared from **(a)** *Tripsacum dactyloides* (2n = 2x =36); and **(b)** *Sorghum bicolor* (2n = 2x = 20). Bars = 10 µm.

Sorghum (*Sorghum bicolor*) and maize diverged from maize ancestors for about 12 MYs (Swigonova et al. 2004). The two maize probes produced background signals on all sorghum chromosomes (**Figure 5b**). Only few of the 10 sorghum chromosomes generated consistent and distinct FISH signal patterns. Thus, the maize probes were not useful to identify individual sorghum chromosomes.

## Discussion

FISH has become the most popular methodology for chromosome identification in plants (Jiang and Gill 2006). Different FISH methodologies have different advantages and drawbacks. BACs and repetitive DNA sequences are the two most common types of FISH probes used in chromosome identification. Unfortunately, BACs are not ideal probes for plant species with large genomes because of the presence of large percentages of repetitive DNA sequences in the clones (Janda et al. 2006; Zhang et al. 2004). Repeat-based chromosome identification systems have been established in several plant species (Findley et al. 2010; Fradkin et al. 2013; Kato et al. 2004; Lengerova et al. 2004; Li et al. 2014; Mukai et al. 1993; Xiong and Pires 2011). Repeat-based systems reveal polymorphisms for the constituent sequences but are not genotype independent. By contrast, single-gene karyotyping probes are genotype independent (Danilova and Birchler 2008; Danilova et al. 2014; Lamb et al. 2007a). Similarly, the FISH signal patterns derived from barcode probes are not polymorphic among different varieties in the same species. A modified pattern would indicate a potential chromosomal rearrangement associated with a cultivar. In addition, barcode probes can also be used for chromosome identification in genetically related species (Braz et al. 2018; Liu et al. 2020) (**Figure 2**). Most importantly, barcode FISH probes can be developed readily in any plant species with a sequenced genome.

We have recently demonstrated that barcode FISH probes developed based on potato sequences were useful to identify individual homoeologous chromosomes from distantly related species, including tomato (*Solanum lycopersicum*), which diverged from potato for 5-8 MYs (Sarkinen et al. 2013; Wang et al. 2008). The potato barcode probes also generated punctuate signals but with a visible background on chromosomes from eggplant (*Solanum melongena*), which diverged from potato for 15.5 MYs (Wu and Tanksley 2010). A total of 54672 oligos were used in the two potato barcode probes, including 16489 oligos (30%) designed from coding sequences of potato genes. In contrast, among the 50,082 maize oligos, only 5525 oligos (11%) are associated with coding sequences. This may explain the fact that the maize barcode probes generate strong background signals on chromosomes of *T. dactyloides*, which diverged from maize for only 4.5 MYs (Hilton and Gaut 1998). Therefore, the percentage of oligos designed from highly conserved DNA sequences is critical to extend the value of barcode FISH probes for chromosome identification in distantly related species.

One of the key factors for developing successful barcode probes is to restrict the oligos within a chromosomal region as narrow as possible. A bright FISH signal can be generated from 1000-2000 oligos. However, these oligos need to be distributed within a few Mb. Each of the 24 regions of our maize barcode span 1.3-3.3 Mb of DNA sequences (**Table S1**). It can be challenging to identify 1000-2000 unique oligos within repetitive chromosomal regions, especially in plant species with very large and complex genomes. Thus, it may become necessary to develop as many oligos as possible within the small islands of genic or single copy sequences in largely repetitive regions. For example, oligos, or overlapping oligos, can be designed from both strands of the DNA sequences within the single copy sequences.

It is not surprising that all *Zea* species and subspecies share a highly conserved karyotype (**Tables 1, 2**) since these species have diverged for only ∼150,000 years (Ross-Ibarra et al. 2009). Phylogenetic studies based on microsatellite markers indicated that *Z. nicaraguensis* and *Z. luxurians* are closely related species and *Z. nicaraguensis* could be treated as a subspecies of *Z. luxurians* (Fukunaga et al. 2005). This conclusion was supported by our barcode FISH and karyotyping data. Terminal knobs, which are distal to the terminal FISH signals, were identified on several chromosome arms, including the long arms of chromosomes 2, 5, 6, 7, 9, and 10 in both *Z. nicaraguensis* and *Z. luxurians* (see also Albert et al. 2010). Interestingly, these terminal knobs were not observed at the same chromosomal positions in other species/subspecies (**Figure 3**). In contrast, subterminal knobs, which are proximal to the terminal FISH signals, were observed on the long arms of chromosomes 2, 5, 6, 7, and 9 (**Figure 3**) in several species/subspecies, but not in *Z. nicaraguensis* and *Z. luxurians*. Thus, *Z. nicaraguensis* and *Z. luxurians* share a similar knob distribution pattern.

A putative paracentric inversion was detected in the long arm of chromosome 4 in both *Z. nicaraguensis* and *Z. luxurians*. Poggio (2005) developed hybrids between *Z. luxurians* and three subspecies of *Z. mays* (*Z. mays ssp. parviglumis, Z. mays ssp. mexicana, Z. mays ssp. mays*) and analyzed meiotic chromosome pairing in hybrids. Interestingly, the most frequent chromosome pairing configuration at metaphase I of meiosis in the hybrids was eight bivalents and four univalents (Poggio et al. 2005), suggesting major structural changes associated potentially with two *Z. luxurians* chromosomes. In addition, bridges and broken chromosome fragments were observed at anaphase I and II (Poggio et al. 2005), which are typical products associated with heterozygous paracentric inversions (Sybenga 1972). In addition, genetic linkage mapping data showed that the chromosome 4 from *Z. nicaraguensis* and *Z. luxurians* share the synteny but they contain a large inversion relative to maize chromosome 4 (Mano and Omori 2013). These results support a paracentric inversion associated with the long arm of chromosome 4. It is likely that additional *Z. nicaraguensis*/*Z. luxurians* chromosomes are involved in major structural changes compared to others in the *Zea* lineage. An inversion was found on the long arm of chromosome 3 based on pachytene chromosome analysis (Fang et al. 2012; Ting 1965), although chromosome 3 shares an identical barcode FISH pattern across all *Zea* species and subspecies (**Figure 3**). Indeed, because the barcode uses only a few well-spaced landmarks per chromosome, chromosomal aberrations, especially those restricted in regions proximal to the landmarks, will go undetected. Barcodes with more dense landmarks and in combination with whole chromosome paints (Albert et al. 2019) will be powerful tools to identify and characterize chromosomal rearrangements occurred during the evolution of *Zea* species.

## Acknowledgements

This research is supported by NSF grant IOS-1444514 and MSU startup funds to J.J.

